# Reversible bacteriophage resistance by shedding the bacterial cell wall

**DOI:** 10.1101/2021.11.17.468999

**Authors:** Veronique Ongenae, Adam Sidi Mabrouk, Marjolein Crooijmans, Daniel Rozen, Ariane Briegel, Dennis Claessen

## Abstract

Phages are highly abundant in the environment and a major threat for bacteria. Therefore, bacteria have evolved sophisticated defense systems to withstand phage attacks. Here, we describe a previously unknown mechanism by which mono- and diderm bacteria survive infection with diverse lytic phages. Phage exposure leads to a rapid and near complete conversion of walled cells to a cell wall-deficient state, which remain viable in osmoprotective conditions and can revert to the walled state. While shedding the cell wall dramatically reduces the number of progeny phages produced by the host, it does not always preclude phage infection. Altogether, these results show that the formation of cell wall-deficient cells prevents complete eradication of the bacterial population and suggest that cell wall-deficiency may limit the efficacy of phage therapy, especially in highly osmotic environments or when used together with antibiotics that target the cell wall.

## Introduction

Bacteria are routinely exposed to a wide range of stresses in their environment, such as changes in temperature, pH, salt concentration or nutrient limitation. To withstand such fluctuating conditions, almost all bacteria are enveloped by a stress-bearing cell wall. Another serious threat are bacteriophages, which outnumber bacteria in the environment by a factor of 10 (*1*). Phages recognize their host by interacting with specific receptors, especially sugars and proteins, present in the cell wall or exposed on the cell surface. Following binding and infection, lytic phages will kill their host, while lysogenic phages persist as prophages inside the bacterium (*2*, *3*). In order to counteract phage attack, bacteria have evolved multiple highly specific escape mechanisms, such as restriction-modification, abortive infection, or CRISPR/Cas systems, among many others (*4*–*13*). Another common strategy to overcome phage attack is by modifying or masking surface associated phage receptors, which is effective, but costly and permanent (*14*–*16*). A better strategy would be one that works for different phages and is reversible, thereby avoiding costs, despite remaining general.

Here, we present a robust response that occurs when lytic phages infect bacterial hosts in osmoprotective environments. We show that upon phage exposure, the bacteria transiently shed their cell wall, leading to the formation of viable cell wall-deficient (CWD) cells that increase survival of the bacterial population.

## Results

### Filamentous actinobacteria can switch to a CWD state after phage infection

Many filamentous actinobacteria have a natural ability to form CWD cells in response to environmental stressors, after which they can revert to filamentous growth upon the removal of stress (*17*–*19*). As part of our work to understand the functions of the transition to CWD cells, we focused on *Streptomyces* strain MBT86, a natural isolate from the Qinling mountains, China (*20*), which has previously been shown to readily produce CWD cells (*18*). When bacteria were grown in osmoprotective conditions (LPB medium) in the presence of the lytic phage LA7 for at least 24 hours, we observed that all remaining cells are CWD after lysis of the dominant mycelial population (Fig. 1A-B, Fig. S1, Movie S1). These cells lacked most of their peptidoglycan-based cell wall and were only enveloped by their cell membrane (Fig. 1C-D). Quantification revealed that the number of CWD cells after 24 hours was more than 34 times higher in the presence of phage LA7 (204 ± 125 CWD cells, n=3) compared to samples without the phage (6 ± 5 CWD cells n=3). When phage LA7 was added to MBT86 in DNB medium (without osmoprotectants), no CWD cells were found (Fig. 1E, Fig. S1A). Consistent with this, the optical density (OD_600_) of MBT86 culture decreased to undetectable levels within 24 hours after addition of the phage, while cultures of MBT86 grown in osmoprotective LPB medium with phage reached significantly higher densities (OD_600_ = 0.285 ± 0.05 after 24 hours). CWD cells were also observed when pre-grown mycelium was exposed to LA7 (Fig. S1C). Viability of CWD cells after phage infection was assessed by determining the number of colony forming units (CFU) after plating mixtures of liquid cultures that still contained the phages. On average, 4.1 x 10^3^ CFU ml^-1^ were observed, which is equivalent to 4% of the population. Furthermore, these colonies consisted of filamentous cells, indicating that the CWD cells had reverted to the canonical mycelial mode-of-growth. Importantly, this demonstrates that a considerable fraction of the population survives phage attack (Fig. S1D-E). Moreover, fewer new phages were produced in osmoprotective LPB medium compared to DNB medium with the same starting concentration of approximately 1000 phages ml^-1^ (Fig. 1F). Taken together, these results show that osmoprotective conditions provide MBT86 with a survival mechanism against phage attack, coinciding with a drop in the number of new progeny phages that are produced.

**Fig. 1.**
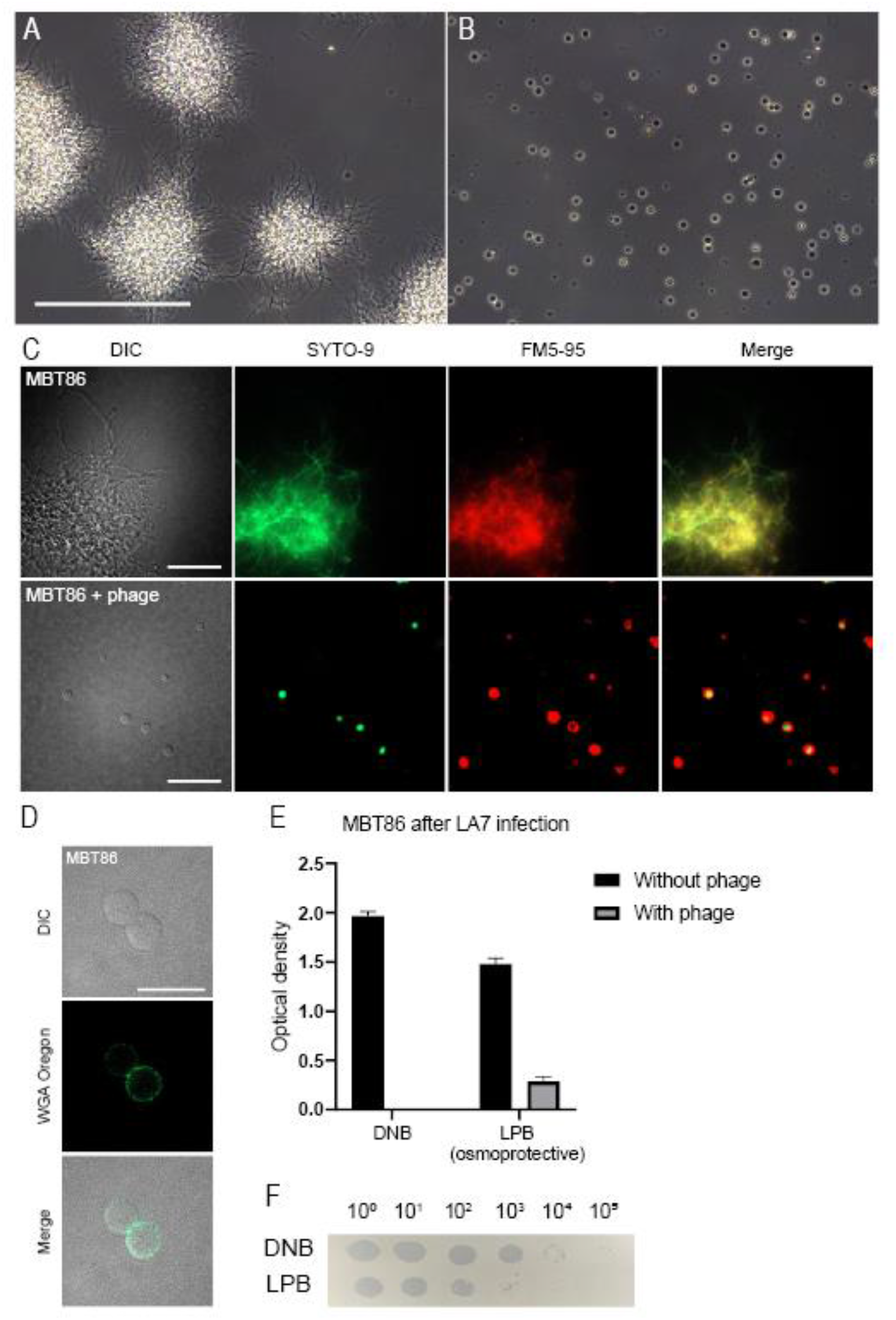
Formation of CWD cells after phage infection in the actinomycete MBT86. (**A**) Morphology of MBT86 after 48 hours in LPB medium. Scale bar represent 100 μm. (**B**) Morphology of MBT86 in LPB medium 24 hours after phage LA7 infection. (**C**) Morphology of MBT86 with and without phage LA7 24 hours after infection. Cells were stained with the DNA dye SYTO-9 (green) and FM5-95 (red) to dye membranes. Scale bars represent 20 μm. (**D**) After 24 hours of phage LA7 infection, the resulting CWD cells of MBT86 were stained with the peptidoglycan stain WGA Oregon. Some cells already start to rebuild their peptidoglycan layer. Scale bar represent 10 μm. (**E**) OD_600_ measurement 24 hours after phage LA7 infection in DNB medium and osmoprotective LPB medium. The experiment was performed in triplicates with the standard deviation presented as error bars. In LPB medium after phage infection, the OD_600_ was still 0.285 ± 0.05. (**F**) Plaque assay showing PFUs of LA7 on DNB and osmoprotective LPB medium. Images were taken 24 hours after LA7 infection.

To determine if the response of MBT86 was specific to LA7, we exposed this strain to three other lytic phages to which MBT86 was susceptible (CE2, CE10 and LD10). In all cases, we observed a qualitatively similar response and a marked increase in CWD cells (Fig. 2A-B). However, exposure to phage LA7 yielded more CWD cells compared to exposure to the other phages CE2, CE10 and LD10. This difference might be explained by a variance of virulence between the phages. A lower population size of LA7 phages was observed after 24 hours (Fig. 2C), which implies that the bacteria probably had more time to grow into longer mycelial filaments that subsequently resulted in more CWD cells. Next, we tested whether other *Streptomyces* strains than MBT86 would also behave similarly to phage exposure. Notably, the number of CWD cells increased after phage infection in all other tested species (Fig. 2D, Fig. S1G).

**Fig. 2.**
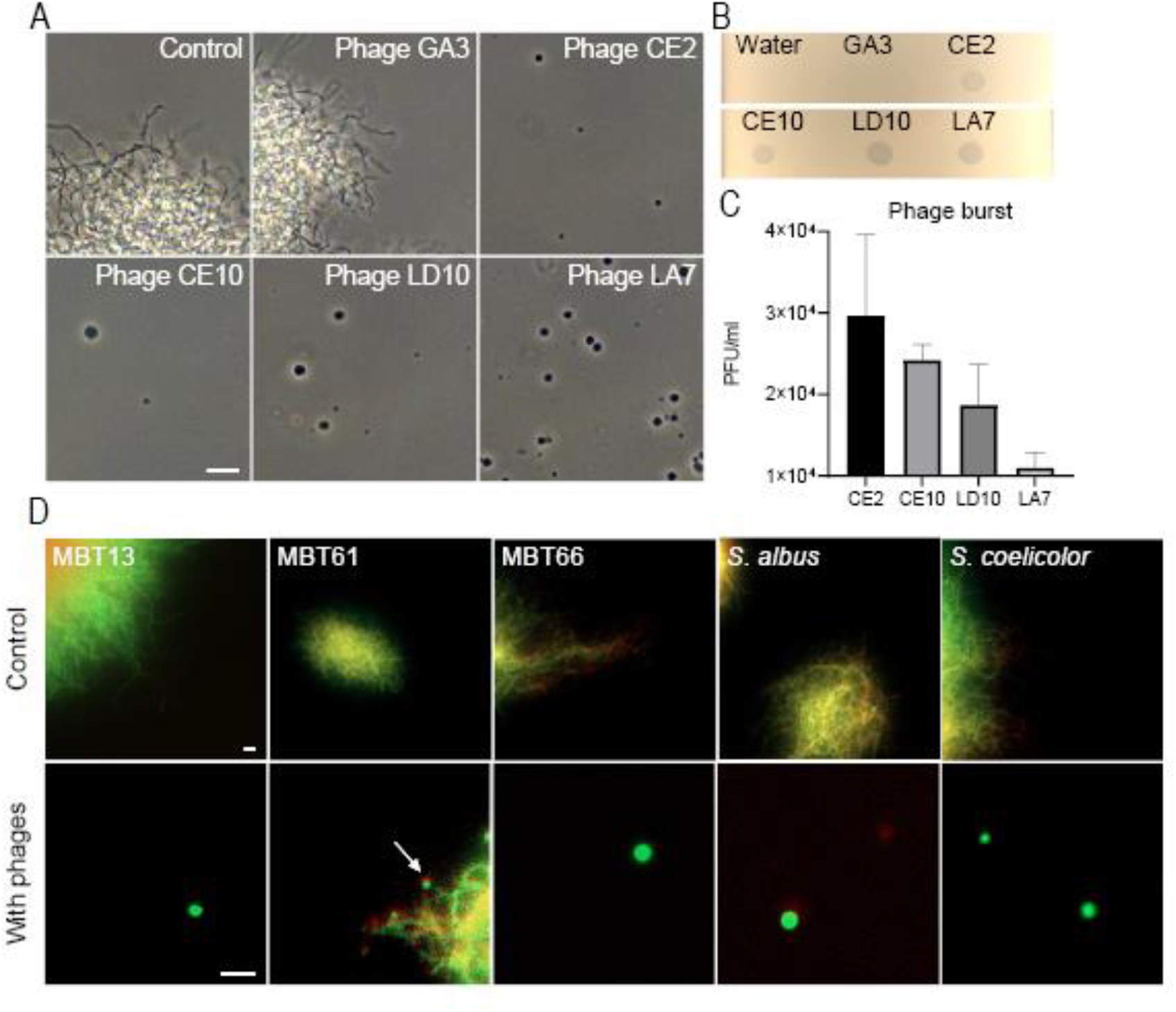
Formation of CWD cells after phage infection is independent of bacteriophage and common in Actinomycetes. (**A**) Morphology of MBT86 24 hours after infection with phages GA3 (non-susceptible), CE2, CE10, LD10 and LA7 in LPB medium. Phages LD10 and LA7 provoke more CWD cell formation compared to phages CE2 and CE10. Scale bar represent 10μm. (**B**) Spot assay of the five phages used in panel A. Plaques indicate susceptibility to the host strain MBT86. (**C**) Phage burst measured 24 hours after MBT86 infection in DNB medium. (**D**). Morphology of CWD cells after phage infection of *Streptomyces albus*, *Streptomyces coelicolor*, and several actinomycetes from our culture collection (referred to with the prefix MBT). Cells were stained with the DNA dye SYTO-9 (green) and FM5-95 (red) to dye membranes. Note that here, the merged image is shown. Scale bars represent 10μm.

### Exposure to phages leads to cell wall-deficiency in Escherichia coli and Bacillus subtilis

Since the formation of CWD cells after phage infection was observed in different *Streptomyces* strains, we wanted to test if this behavior is also found in other genera. Therefore, we extended our studies to the phylogenetically distant *Bacillus subtilis* and *Escherichia coli*, which split from *Streptomyces* roughly three billion years ago (*21*). Both species were inoculated in osmoprotective LPB medium and exposed to the lytic phages φ29 and T4, respectively. 24 hours after the addition of the phage, both cultures contained CWD cells, which did not appear to proliferate (Fig. 3). To quantify CWD cell formation after phage infection in soft agar, samples with and without the addition of phages were analyzed. To exclude rod-shaped bacteria, only particles with a circularity of 0.8-1.00 were included in the analysis. We observed that all *B. subtilis* cells embedded in soft agar (another osmoprotective environment) adopted a CWD lifestyle approximately 12 hours after φ29 phage infection (Fig. 4A, Movie S2). Without addition of the phage, no CWD cells were observed. A comparable response was observed with *E. coli*, where almost 93% of the surviving cells had adopted a wall-deficient state after 24 hours (Fig. 5A, Movie S3). To explore whether the formation of CWD cells was due to endolysins extruded by lysed bacteria, a phage infected *B. subtilis* culture in LPB was filtered and only the supernatant was added to a fresh bacterial culture in LPB. However, this did not give rise to CWD cells, suggesting there are insufficient free lytic agents secreted in the medium to induce this response.

**Fig. 3.**
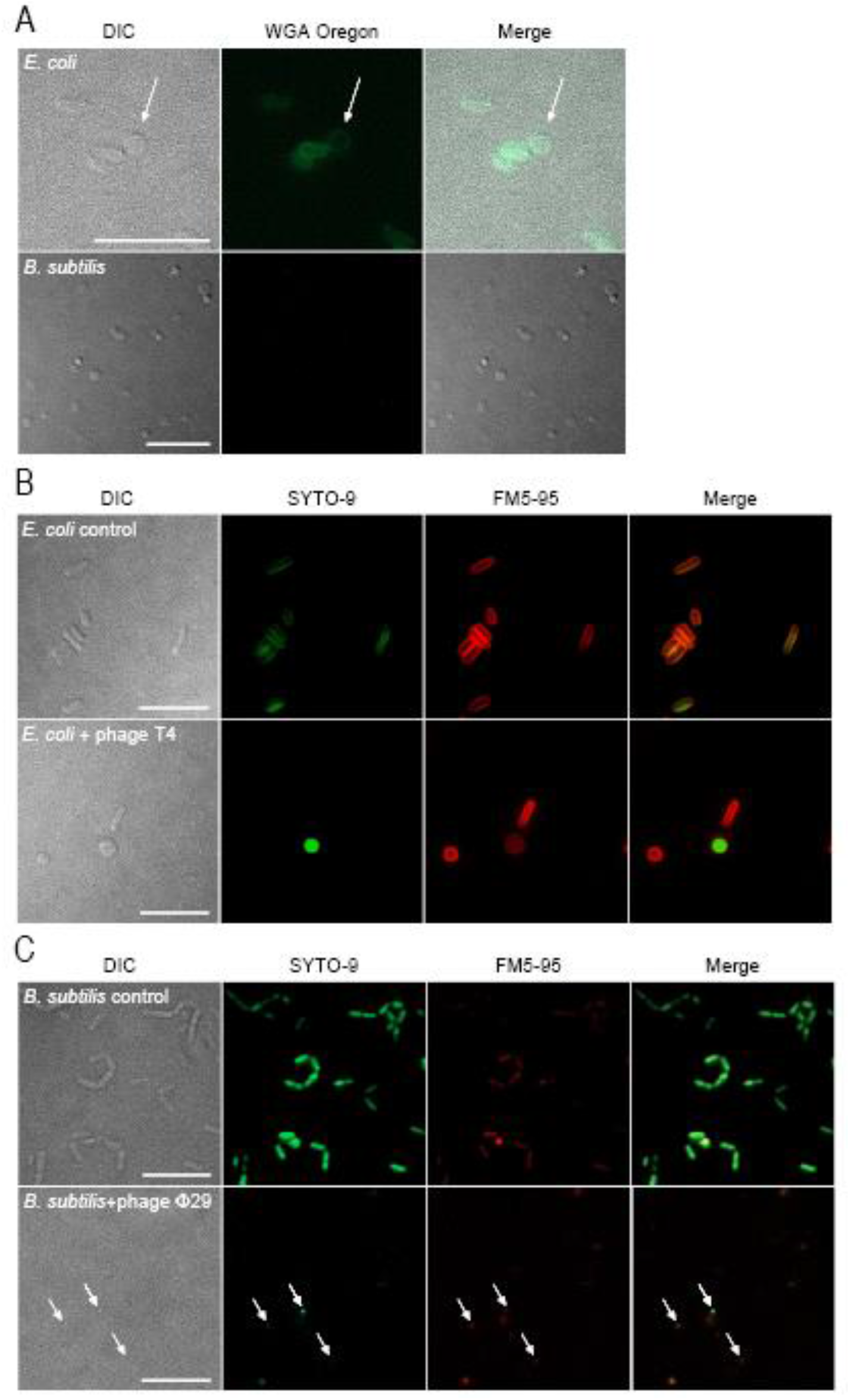
Formation of CWD cells after phage infection of *E. coli* and *B. subtilis* in LPB medium. (**A**) *E. coli* and *B. subtilis* were inoculated in LPB medium for 24 hours with phages T4 and φ29, respectively. CWD cells were stained with WGA Oregon to visualize peptidoglycan. Note that *E. coli* CWD cells still have some peptidoglycan, while *B. subtilis* cells have none. (**B**) Morphology of *E. coli* with and without phage T4 after 24 hours in LPB medium. Cells were stained with the DNA dye SYTO-9 (green) and FM5-95 (red) to dye membranes. Scale bars represent 10μm. (**C)** Morphology of *B. subtilis* with and without phage φ29 after 24 hours in LPB medium. Cells were stained with the DNA dye SYTO-9 (green) and FM5-95 (red) to dye membranes. Scale bars represent 10μm. Note that CWD cells from *B. subtilis* are smaller than CWD cells from *E. coli* and MBT86.

**Fig. 4.**
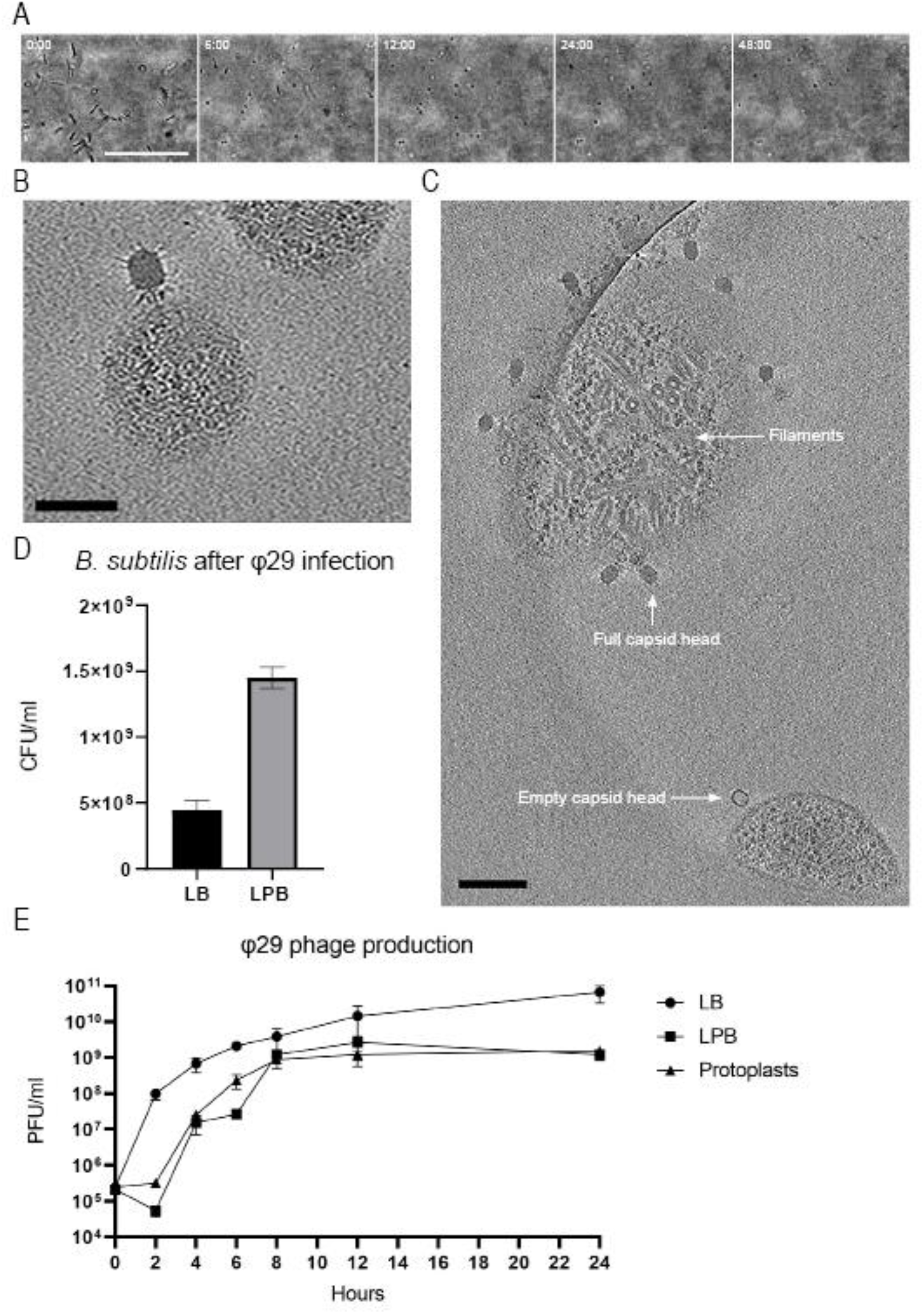
Growth, cryo-ET and infection rate of *B. subtilis* after φ29 phage infection. (**A**) Time-lapse of *B. subtilis* after φ29 phage infection followed for 48 hours. Individual micrographs were taken from supplementary movie 1. Scale bars represent 50μm. (**B**) Phage φ29 attaches to the membrane of a CWD *B. subtilis* cell. Scalebar represents 50nm. (**C**) A tomography snapshot of *B. subtilis* protoplasts with φ29, see supplementary movie 3 for the full tomography. On the bottom is an empty phage attaching to a protoplast, while on top there is a protoplast filled with filaments associated with phage φ29 DNA assembly. Scalebar represents 100nm. (**D**) CFU’s of *B. subtilis* 24 hours after phage infection with φ29 in LB medium and LPB medium. Significantly more cells survive in LPB medium (P < 0.0001). Phage φ29 production measured in plaque forming units (PFU)/ml in LB and LPB media with protoplasts as a negative control. The φ29 production in LB media compared to LPB media is significantly different (P < 0.027) at T=24. Statistics was performed with a Student’s T-Test.

**Fig. 5.**
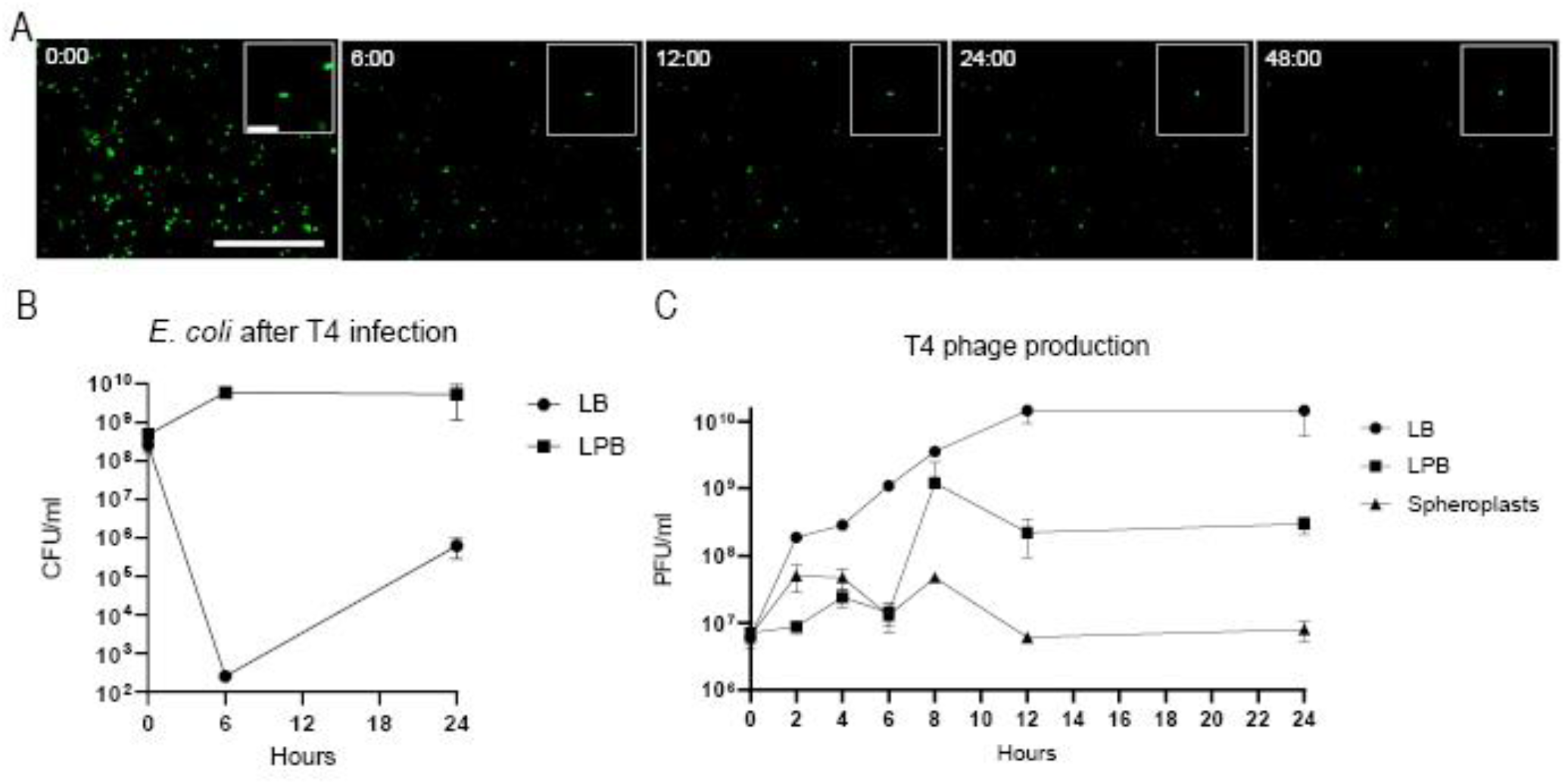
Growth and infection rate of *E. coli* after T4 phage infection. (**A**) Time-lapse of green fluorescence *E. coli* after T4 phage infection followed for 48 hours. Individual micrographs were taken from supplementary movie 1. Scale bars represent 100μm and 10μm for the inlays. (**B**) CFU’s of *E. coli* 24 over time after phage infection with T4 in LB medium and LPB medium. More cells survive after 24 hours in LPB medium (P = 0.09). (**C**) T4 phage production measured in PFU ml^-1^ in LB and LPB media with spheroplasts as a negative control. The T4 production in LB media compared to LPB media is significantly different (P < 0.05) for each timepoint after T=0. All statistics was performed with a Student’s T-Test.

### Artificially produced CWD cells survive phage infection

To test whether artificially produced CWD cells were susceptible to phage infection, the cell wall was stripped away using lysozyme and/or antibiotics. These cells are referred to as protoplasts for monoderm bacteria and spheroplasts for diderm bacteria and do not proliferate without their cell wall. Protoplasts of MBT86 and *B. subtilis* as well as spheroplasts of *E. coli* were seemingly unaffected by phage infection and did not disappear from the culture after exposure to phages LA7, φ29 and T4, respectively (Fig. S2). These results provide direct evidence that wall-deficiency provides protection against lytic phages. Interestingly, phages were found near artificially produced protoplasts and spheroplasts of *B. subtilis* and *E. coli* (Fig. S3). This raised the question whether the phages could attach and maybe even eject their genomic material into wall-deficient cells.

### Phages cannot attach to CWD cells of E. coli, but can attach and propagate in B. subtilis CWD cells

To address this question, we used cryo-electron tomography (cryo-ET). The resulting three-dimensional data of *E. coli* spheroplasts with phage T4 revealed a highly heterogeneous sample with an inconsistent appearance of CWD cells and outer membrane vesicles. However, no phage attachment was observed for any of the CWD cells or cell fragments that were imaged (Fig. S4, Movie S4). Interestingly, cryo-ET data of *B. subtilis* showed that bacteriophage φ29 was not only able to attach to protoplasts of *B. subtilis* (Fig. 4B) but could also eject DNA into the cells and propagate. We observed attached phages with filled as well as empty capsid heads (indicating DNA ejection). Furthermore, we detected filaments inside some protoplasts that were surrounded by phages (Fig. 4C, Fig. S5, Movie S5). These filaments strongly resemble p1 proteins that are associated with φ29 DNA replication (*22*).

In both *B. subtilis* and *E. coli*, significantly more bacteria survived a phage infection in the osmoprotective LPB medium compared to LB medium (Fig. 4D, Fig. 5B). In addition, *B. subtilis* grown in LPB medium and pre-made protoplasts produced similar amounts of φ29 progeny, in contrast to LB medium, where a significantly higher phage titer was observed over time (Fig. 4E). When the same experiment was performed with *E. coli*, we again noticed increased levels of phage T4 in LB medium and significantly fewer T4 progeny in LPB medium over time, as opposed to spheroplasts, that did not produce any progeny at all (Fig. 5C). These results are in line with our cryo-ET data showing that phage φ29 was able to attach and propagate in protoplasts of *B. subtilis*, while phage T4 was not observed in association with any *E. coli* spheroplasts remnants.

Altogether, our results imply that transiently lacking or shedding a cell wall is a common and general escape mechanism against phage attack.

## Discussion

The arms-race between bacteria and bacteriophages has resulted in the evolution of many antiphage-defense mechanisms, which we recently started to uncover (*23*, *24*). In this study, we have discovered a yet unknown general escape mechanism against phage attack for both the diderm model organism *E. coli*, the monoderm *B. subtilis*, and various filamentous actinomycetes. In contrast to L-forms, the CWD cells produced after phage infection are not able to proliferate and will switch back to the walled mode-of-growth in favorable conditions (*18*, *25*). Besides the species tested here, many more bacteria may be capable of surviving phage infection in osmoprotective medium. In standard hypotonic cultivation media without high levels of osmolytes, phages produce endolysins that degrade the bacterial cell wall and cause lysis. For *B. subtilis* it has been shown that prophage-encoded endolysins in LB medium resulted in the release of membrane vesicles that immediately burst (*26*). However, our results show that the addition of spent media from cultures exposed to phages did not result in the formation of CWD cells in LPB medium. We hypothesize that the concentration of endolysins in the supernatant is either not high enough to initiate the conversion to CWD cells, or these wall-deficient cells are simply a result of the initial attachment or DNA ejection of phages in supportive medium. In nature, bacteria may encounter such osmoprotective environments, for example, in eukaryotic host cells, plant sap with high levels of osmolytes or in soil after rain (*27*–*31*).

Cryo-ET allowed us to observe the distinct interaction between bacteriophages and CWD cells. To our knowledge, this is the first time the interaction of CWD cells and bacteriophages was captured in detail. We showed that φ29 was able to attach, eject DNA and even produce new progeny phages in *B. subtilis* protoplasts, as indicated by filaments inside protoplasts (*22*). Phage φ29 normally binds to glycosylated teichoic acid in the cell wall (*32*), but can sporadically also attach to fragments of lipoteichoic acids in the membrane of protoplasts (*33*, *34*). For CWD *E. coli* cells, it was hypothesized that T4 could recognize OmpC and lipopolysaccharide on the remaining outer membrane (*35*). However, three-dimensional cryo-ET imaging revealed that T4 phages were not attached to remnants of *E. coli* spheroplasts. On the other hand, a recent paper from Petrovic Fabijan *et al*., showed that T4 was able to eject DNA in L-forms of *E. coli*, but no cells were lysed (*36*). Perhaps, the difference in attachment of phages could be explained by the fact that L-forms are able to proliferate, whereas spheroplasts cannot. However, no new progeny phages were produced in either L-forms or spheroplasts of *E. coli* and we both observed an increased survivability in such CWD cells. Not only *E. coli*, but also MBT86 and *B. subtilis* have a significantly higher survival after phage attack in osmoprotective medium by adopting a CWD state, compared to walled cells in normal medium.

The ability to shed the cell wall after phage infection could have major consequences that limits the use of phage therapy. Phage therapy is considered a promising alternative to treat antibiotic resistant infections (*29*). However, in this study we have shown that both diderm and monoderm bacteria can escape phage attack by adopting a cell wall-deficient lifestyle. These CWD cells can arise in osmoprotective environments in nature, or in the presence of cell wall targeting antibiotics. Whereas commonly known phage defense mechanisms like restriction modification or CRISPR/Cas can protect bacteria against specific phages, CWD cells might be resistant to a broad range of phages, which warrants a careful analysis in the interaction between phages and their host.

## Methods

### Strains, bacteriophage isolation and sequencing

Bacterial strains and phages used in this work are listed in Supplementary Table 1. Bacteriophages were isolated from soil samples collected in the Netherlands at longitude N52°23’31” and latitude E4°34’49”. The soil samples were collected at a depth of three cm and stored at 4°C before processing. Actinobacteriophage isolation was performed as described before using *Streptomyces lividans* (phage LA7 and LD10)*, Streptomyces coelicolor* (phage CE2 and CE10) and *Streptomyces griseus* (phage GA3) as bacterial hosts in Difco Nutrient Broth (DNB) (*37*). High viral titer stocks were obtained by picking streak purified plaques until confluent lysis was observed. Plates were flooded with DNB broth and lysates were filtered through a 0.22μm filter. Concentration and activity of phages was confirmed using a spot assay. Phages were serial diluted and 3μl was spotted on a DNB double agar overlay plate containing the host strain. After 24 hours, PFU’s were quantified. Genomic material of bacteriophage LA7 was isolated using the Phage DNA Isolation Kit (Norgen) according to the manufacturers protocol. Whole-genome sequencing of phage LA7 followed by *de novo* assembly was performed by BaseClear (Leiden, The Netherlands).

### Characterization of bacteria and phage interactions

L-phase broth (LPB) was used to identify if Streptomyces, *E. coli*, and *B. subtilis* were able to form CWD cells after phage infection. LPB is an osmoprotective medium, which consists of a mixture of TSBS and YEME (1:1 v/v) supplemented with 25mM MgCl_2_ (*18*). For Streptomyces strains, 10^6^ spores ml^-1^ were inoculated in 10 ml LPB medium with 1000 phages ml^-1^. OD_600_ was measured after 24 hours of incubation at 30°C, 200 RPM. To test if *E. coli* and *B. subtilis* could form CWD cells, an overnight culture was diluted to OD = 0.01 in LPB and incubated at 30°C until OD = 0.5 was reached. Subsequently, phages T4 or φ29 was added at a MOI of 2.0. The culture was incubated for 24 hours at 30°C, 200 RPM before imaging.

To produce protoplasts, MBT86 was grown in a mixture of TSBS and YEME (1:1 v/v) supplemented with 5mM MgCl_2_ and 0.5% glycine for 48 hours at 30°C, 100 RPM. Protoplasts were prepared by incubating the mycelial pellet in 10mg/ml lysozyme solution for one hour (*38*). The culture was filtered through an EcoCloth™ filter, centrifuged at 1000g for seven minutes and resuspended in one ml P+ buffer (*38*). Protoplasts of *B. subtilis* were produced using an overnight culture that was diluted to OD=0.01 in LPB medium supplemented with 25 mM MgCl_2_, 1.2 mg/ml penicillin G sodium (Duchefa Biochemie) and 8 mg/ml lysozyme, and grown for 24 hours at 30°C. For *E. coli* spheroplasts, an overnight culture was diluted to OD = 0.01 in LPB medium supplemented with 25mM MgCl_2_ and 0.4 mg/ml penicillin G sodium, and was grown for 24 hours at 30°C. Subsequently, MBT86 protoplasts were inoculated with 1000 phages ml^-1^, while *E. coli* spheroplasts and *B. subtilis* protoplasts were inoculated with phages T4 or φ29 at a MOI of 1.0 and incubated for 24 hours at 30°C, 200 RPM before imaging.

To determine if proteins extruded after phage infection can initiate the formation of CWD cells, an overnight *B. subtilis* culture was inoculated in LPB medium until OD = 0.5 and subsequently infected with φ29 at a MOI of 10. The medium was filtered after 24 hours using an Amicon ultra-15 centrifugal filter unit (30kDa, Millipore) and the supernatant was added to a fresh overnight grown *B. subtilis* culture at OD = 0.5. After 24 hours, CWD cell formation was assessed.

### Viability after phage infection

To verify the viability of the CWD cells from MBT86 observed after phage infection, 100μl of the phage-infected culture in DNB or LPB medium was plated in triplicate on MYM medium (*39*). The plates were grown for 3 days at 30°C before analyzing morphology. CFU’s of *E. coli* and *B. subtilis* were determined using serial dilution of the phage infected culture on LB plates. After one day of growth at 30°C, CFU’s were calculated. To calculate the concentration of phages after infection, 1ml of a phage-infected culture (triplicates) was filtered through 0.22μm, serial diluted and PFU’s were quantified with a spot assay.

### Microscopy

The Zeiss Axio Lab A1 upright Microscope, equipped with an Axiocam 105 color (resolution of 5 megapixel) or Axiocam MRc (resolution of 64.5 nm/pixel) camera was used to take bright field images. The Zeiss Axio observer Z1 microscope was used to visualize stained bacteria on a thin layer of LPB soft agar covering the glass slide. All stains (Molecular Probes™) were added directly to 25μl of liquid culture. The membrane dye FM5-95 was used at a final concentration of 0.02 mg/ml and the nucleic acid dye SYTO-9 at 0.5μM. Peptidoglycan was visualized using 0.02mg/ml Wheat Germ Agglutinin (WGA) Oregon. SYTO-9 and WGA Oregon were excited using a 488nm laser and monitored in the region between 505-600nm, while FM5-95 was excited at 561nm and detected using a 595nm long pass filter. Images were adjusted using OMERO software (*40*).

Fluorescent labeling of T4 and φ29 phages was accomplished by adding 475μl phage with 25μl Alexa Fluor™ 488 NHS-Ester (Thermo Fisher) at a final concentration of 0.05 mg/ml (10mM). The labeled phages were gently shaked at room temperature for 3 hours and washed with an Amicon ultra-0.5 centrifugal filter unit (100kD, Millipore) in phage storage buffer (100mM NaCl, 10mM MgSO_4_, 10mM Tris-HCl and 1mM EDTA) and stored at 4°C until use.

The OrganoPlate^®^ 2-lane 96 wells plate was used to visualize phage-host interaction over time. An overnight grown culture of MBT86, *E. coli* and *B. subtilis* was inoculated in 0.5% DNB or LB soft agar, while >10^6^ phages ml^-1^ were inoculated on the other side of the PhaseGuide™ to allow free interaction between the channels. Time-lapses were recorded using the Lionheart™ FX Automated Microscope with temperature set at 30°C for at least 48 hours. Images were analyzed using Gen5™ software.

### Cryo-electron microscopy

*E. coli* spheroplasts and *B. subtilis* protoplasts were produced as described before. After 24 hours, the cultures were centrifuged at 300 RPM for 30 min and the pellet was resuspended with 10 nm gold fiducial markers (Cell Microscopy Core, UMC Utrecht) at a 1:10 ratio. After 10 min of incubation approximately 10^8^ to 10^10^ phages were added to visualize initial interaction. Of this sample, 3μl was added to a glow-discharged R2/2 200 mesh holey carbon EM grid (Quantifoil) for 30 seconds and the sample was blotted for 1 second, after which the grid was plunged in liquid ethane using the Leica GP automated freeze-plunger. The vitrified samples were observed at the Netherlands Center for Nanoscopy (NeCEN, Leiden). Tilt series of *E. coli* were collected with the 300 kV Titan Krios 2 TEM (Thermo Fisher Scientific) using a X-FEG electron gun and Gatan K2 Summit DED (Gatan Pleasanton). The tilt series ranged from −60 to +60 with a two-degree increment, had a defocus of 8, a pixel size of 4.4 Å and a cumulative dose of 100 electrons per Å^2^. Tilt series of *B. subtilis* were collected with the 300 kV Titan Krios 1 TEM (Thermo Fisher Scientific) using a S-FEG electron gun and K3 bioquantum (Gatan Pleasanton). The tilt series settings are the same as for *E. coli*. Tomography software (Thermo Fisher Scientific) was used to record the data and IMOD software (*41*) to reconstruct tomograms.

### Data and materials availability

All data are available in the main text and supplementary materials. Phage genome sequencing data are available on NCBI (GenBank Accession No. OK412919).

## Acknowledgments

We would like to thank Dr. J. J. Willemse for the help with analyzing images on ImageJ (FIJI) and Dr. W. Noteborn for assisting and obtaining cryo-ET data. We are also thankful to Eline Duits for isolating *Streptomyces* phages. and Dr. Franklin Nobrega for sending us phage φ29.

## Author contributions

Conceptualization: VO, AB, DC

Investigation: VO, AS, MC

Visualization: VO

Funding acquisition: AB, DC

Writing – Original draft: VO

Writing – Review & Editing: VO, DR, AB, DC

## References

1. G. P. C. Salmond, P. C. Fineran, in Nature Reviews Microbiology. (Nature Publishing Group, 2015), vol. 13, pp. 777–786.

2. A. Das, S. Mandal, V. Hemmadi, V. Ratre, M. Biswas, Studies on the gene regulation involved in the lytic-lysogenic switch in *Staphylococcus aureus* temperate bacteriophage Phi11. The Journal of Biochemistry 168, 659–668 (2020).

3. R. Edgar et al., Bacteriophage infection is targeted to cellular poles. Molecular Microbiology 68, 1107–1116 (2008).

4. R. L. Dy, R. Przybilski, K. Semeijn, G. P. C. Salmond, P. C. Fineran, A widespread bacteriophage abortive infection system functions through a type IV toxin-antitoxin mechanism. Nucleic Acids Research 42, 4590–4605 (2014).

5. A. Harms, D. E. Brodersen, N. Mitarai, K. Gerdes, in Molecular Cell. (Cell Press, 2018), vol. 70, pp. 768–784.

6. F. Hille et al., in Cell. (Cell Press, 2018), vol. 172, pp. 1239–1259.

7. C. M. Johnson, M. M. Harden, A. D. Grossman, in bioRxiv. (bioRxiv, 2020), pp. 2020.2012.2013.422588.

8. A. Lopatina, N. Tal, R. Sorek, in Annual Review of Virology. (Annual Reviews Inc., 2020), vol. 7, pp. 371–384.

9. K. S. Makarova et al., in Nature Reviews Microbiology. (Nature Research, 2020), vol. 18, pp. 67–83.

10. L. M. Malone, N. Birkholz, P. C. Fineran, in Current Opinion in Biotechnology. (Elsevier Ltd, 2021), vol. 68, pp. 30–36.

11. Y. Qiu, S. Wang, Z. Chen, Y. Guo, Y. Song, An active type I-E CRISPR-cas system identified in *Streptomyces avermitilis*. PLoS ONE 11, (2016).

12. M. R. Tock, D. T. F. Dryden, in Current Opinion in Microbiology. (Elsevier Current Trends, 2005), vol. 8, pp. 466–472.

13. X. Wu, J. Zhu, P. Tao, V. B. Rao, Bacteriophage T4 escapes CRISPR attack by minihomology recombination and repair. mBio, (2021).

14. H. Harvey et al., *Pseudomonas aeruginosa* defends against phages through type IV pilus glycosylation. Nature Microbiology 3, 47–52 (2018).

15. J. T. Rostøl, L. Marraffini, in Cell Host and Microbe. (Cell Press, 2019), vol. 25, pp. 184–194.

16. D. Scholl, S. Adhya, C. Merril, *Escherichia coli* K1’s capsule is a barrier to bacteriophage T7. Applied and Environmental Microbiology 71, 4872–4874 (2005).

17. A. Du Toit, Living without the cell wall. Nature Reviews Microbiology 17, 65 (2019).

18. K. Ramijan et al., Stress-induced formation of cell wall-deficient cells in filamentous actinomycetes. Nature Communications 9, (2018).

19. E. Ultee, K. Ramijan, R. T. Dame, A. Briegel, D. Claessen, Stress-induced adaptive morphogenesis in bacteria. Advances in Microbial Physiology 74, 97-141 (2019).

20. H. Zhu et al., Eliciting antibiotics active against the ESKAPE pathogens in a collection of actinomycetes isolated from mountain soils. Microbiology (United Kingdom) 160, 1714–1726 (2014).

21. F. U. Battistuzzi, A. Feijao, S. B. Hedges, A genomic timescale of prokaryote evolution: insights into the origin of methanogenesis, phototrophy, and the colonization of land. BMC evolutionary biology 4, 1–14 (2004).

22. A. Bravo, M. Salas, Polymerization of bacteriophage ø29 replication protein p1 into protofilament sheets. The EMBO journal 17, 6096–6105 (1998).

23. A. B. Isaev, O. S. Musharova, K. V. Severinov, Microbial arsenal of antiviral defenses. part II. Biochemistry (Moscow) 2021 86:4 86, 449–470 (2021).

24. A. B. Isaev, O. S. Musharova, K. V. Severinov, in Biochemistry (Moscow). (Pleiades journals, 2021), vol. 86, pp. 319–337.

25. J. Errington, L-form bacteria, cell walls and the origins of life. Open Biology 3, (2013).

26. M. Toyofuku et al., Prophage-triggered membrane vesicle formation through peptidoglycan damage in *Bacillus subtilis*. Nature Communications 8, 1–10 (2017).

27. F. Germano, D. Testi, L. Campagnolo, M. Scimeca, C. Arcuri, in medRxiv. (medRxiv, 2020), pp. 2020.2007.2013.20120428.

28. V. Grosboillot, Revertant *Listeria monocytogenes* and conditions for their persistence as intracellular L-forms. (2021).

29. K. M. Mickiewicz et al., Possible role of L-form switching in recurrent urinary tract infection. Nature Communications 10, (2019).

30. A. M. Paton, C. M. J. Innes, Methods for the establishment of intracellular associations of L-forms with higher plants. Journal of Applied Bacteriology 71, 59–64 (1991).

31. J. M. Wood, Bacterial responses to osmotic challenges. The Journal of General Physiology 145, 381 (2015).

32. F. Young, Requirement of glucosylated teichoic acid for adsorption of phage in *Bacillus subtilis* 168. Proceedings of the National Academy of Sciences of the United States of America 58, 2377 (1967).

33. M. M. Farley, J. Tu, D. B. Kearns, I. J. Molineux, J. Liu, Ultrastructural analysis of bacteriophage Φ29 during infection of *Bacillus subtilis*. Journal of structural biology 197, 163 (2017).

34. E. D. Jacobson, O. E. Landman, Adsorption of bacteriophages phi 29 and 22a to protoplasts of *Bacillus subtilis* 168. Journal of Virology 21, 1223–1227 (1977).

35. A. Washizaki, T. Yonesaki, Y. Otsuka, Characterization of the interactions between *Escherichia coli* receptors, LPS and OmpC, and bacteriophage T4 long tail fibers. MicrobiologyOpen 5, 1003–1015 (2016).

36. A. Petrovic Fabijan et al., L-form switching confers antibiotic, phage and stress tolerance in pathogenic *Escherichia coli*. bioRxiv, 2021.2006.2021.449206 (2021).

37. J. E. Dowding, Characterization of a bacteriophage virulent for *Streptomyces coelicolor* A3(2). Journal of General Microbiology 76, 163–176 (1973).

38. T. Kieser, M. J. Bibb, M. J. Buttner, K. F. Chater, D. A. Hopwood, Practical streptomyces genetics. 291, (2000).

39. C. Stuttard, Temperate phages of *Streptomyces venezuelae*: lysogeny and host specificity shown by phages SV1 and SV2. Journal of General Microbiology 128, 115–121 (1982).

40. C. Allan et al., OMERO: flexible, model-driven data management for experimental biology. Nat Methods 9, 245–253 (2012).

41. J. R. Kremer, D. N. Mastronarde, J. R. McIntosh, Computer visualization of three-dimensional image data using IMOD. Journal of structural biology 116, 71–76 (1996).

