## Supplementary Dataset for "Reversible bacteriophage resistance by shedding the bacterial cell wall"

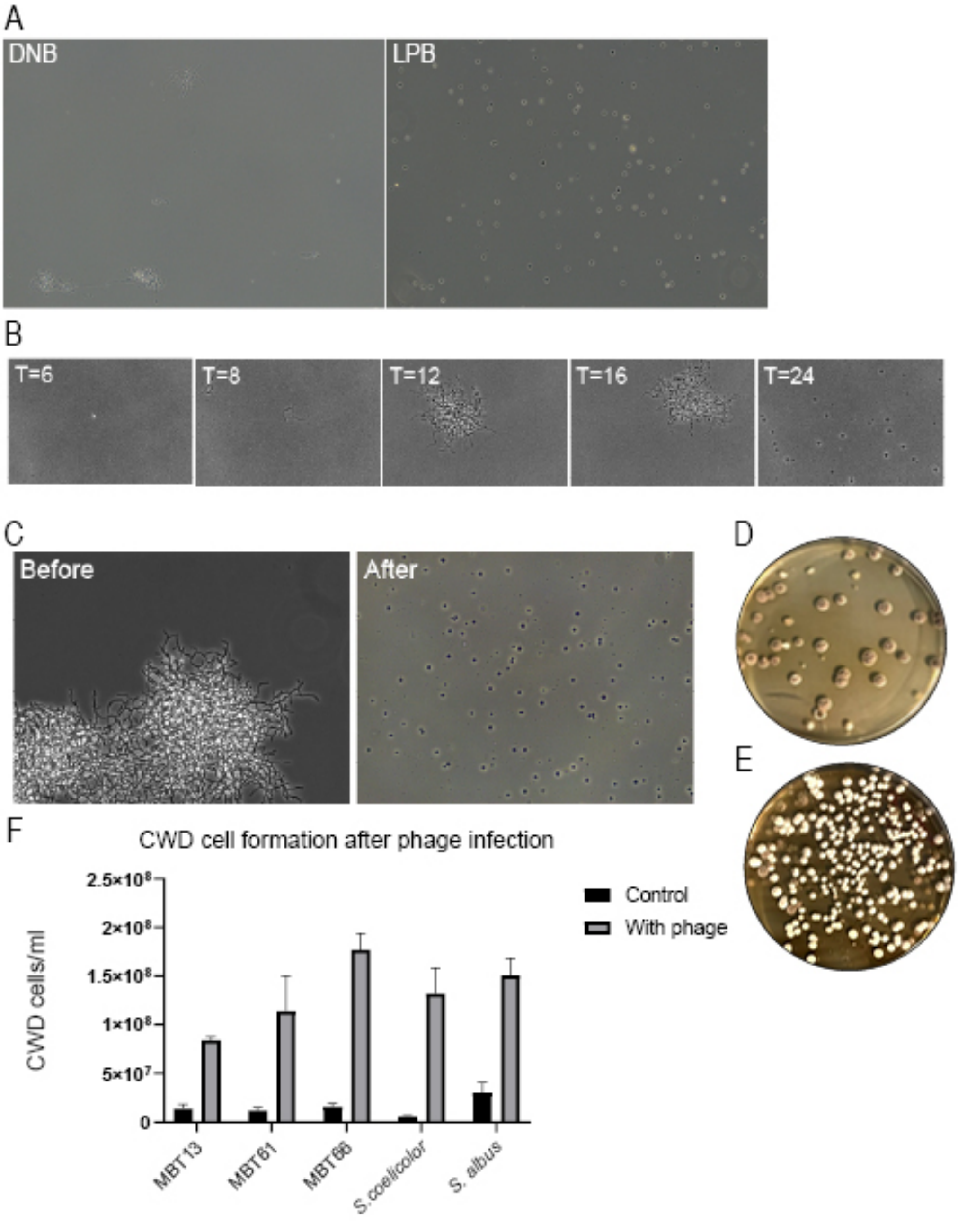


**Fig. S1. Morphology of actinomyces after phage infection.** (**A**) Morphology of MBT86 after 24 hours of phage infection with LA7 in DNB medium and LPB medium. (**B**) MBT86 spores and phage LA7 were inoculated simultaneously in LPB medium at T=0. After approximately 8 hours, the spores start to form mycelial fragments that turned into CWD cells after 24 hours. (**C**) MBT86 spores were pre-grown for 24 hours to form mycelium in LPB before phage LA7 was added. After another 24 hours, only CWD cells were observed. (**D**) Regrowth of MBT86 after phage infection in DNB medium. Note that small mycelial fragments as seen in panel A grew back into colonies. (**E**) Regrowth of MBT86 CWD cells after phage infection in LPB medium. Note that these colonies have developmental defects, as seen by a mixture of white and grey colonies. (**F**) CWD cells were counted with an hemacytometer in duplo of several Streptomyces strains in LPB with and without phages.

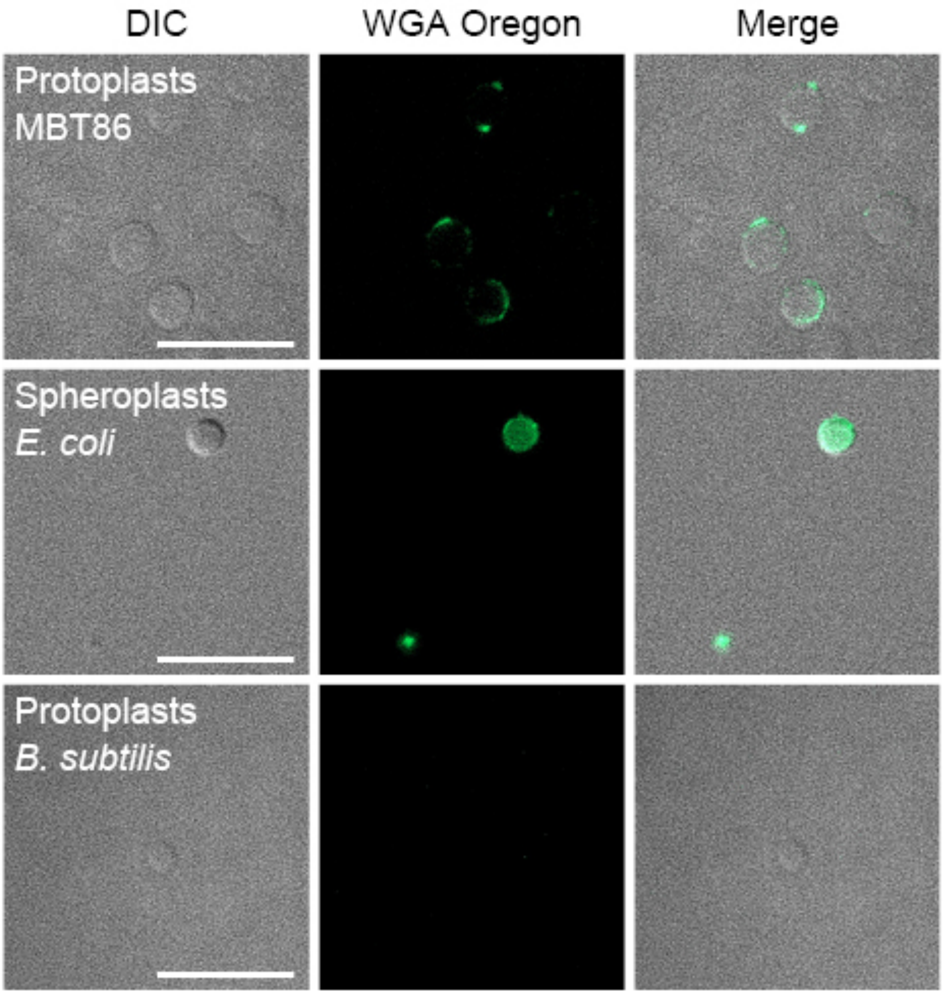


**Fig. S2. Artificially produced CWD cells survive phage infection**. MBT86 protoplasts (first panel), *E. coli* spheroplasts (second panel) and *B. subtilis* protoplasts (third panel) were grown in LPB medium with either phage LA7, T4 or φ29 and imaged after 24 hours. Cells were stained with WGA Oregon to visualized peptidoglycan. Protoplasts of *B. subtilis* have no peptidoglycan, similar to *B. subtilis* CWD cells formed after phage infection. Scale bars represents 10 µm.

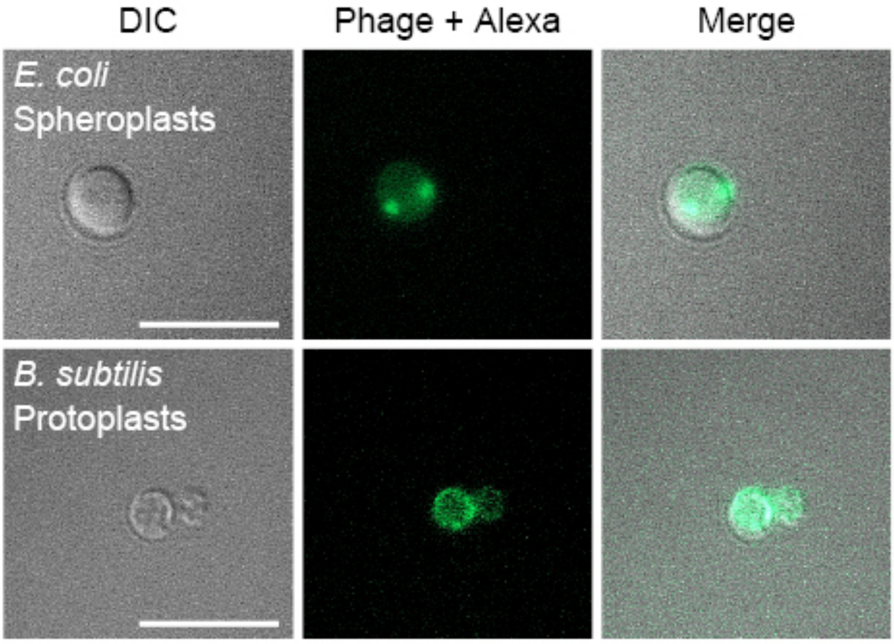


**Fig. S3. Phages are observed in close proximity to artificially produced CWD cells**. *E. coli* strain ET8 spheroplasts and *B. subtilis* protoplasts were imaged 24 hours after inoculation with phages T4 or φ29 that were previously dyed with Alexa Fluor™ 488 NHS-Ester. The CWD cells did not lyse and phages surrounded the cells. Scale bars represents 10 μm.

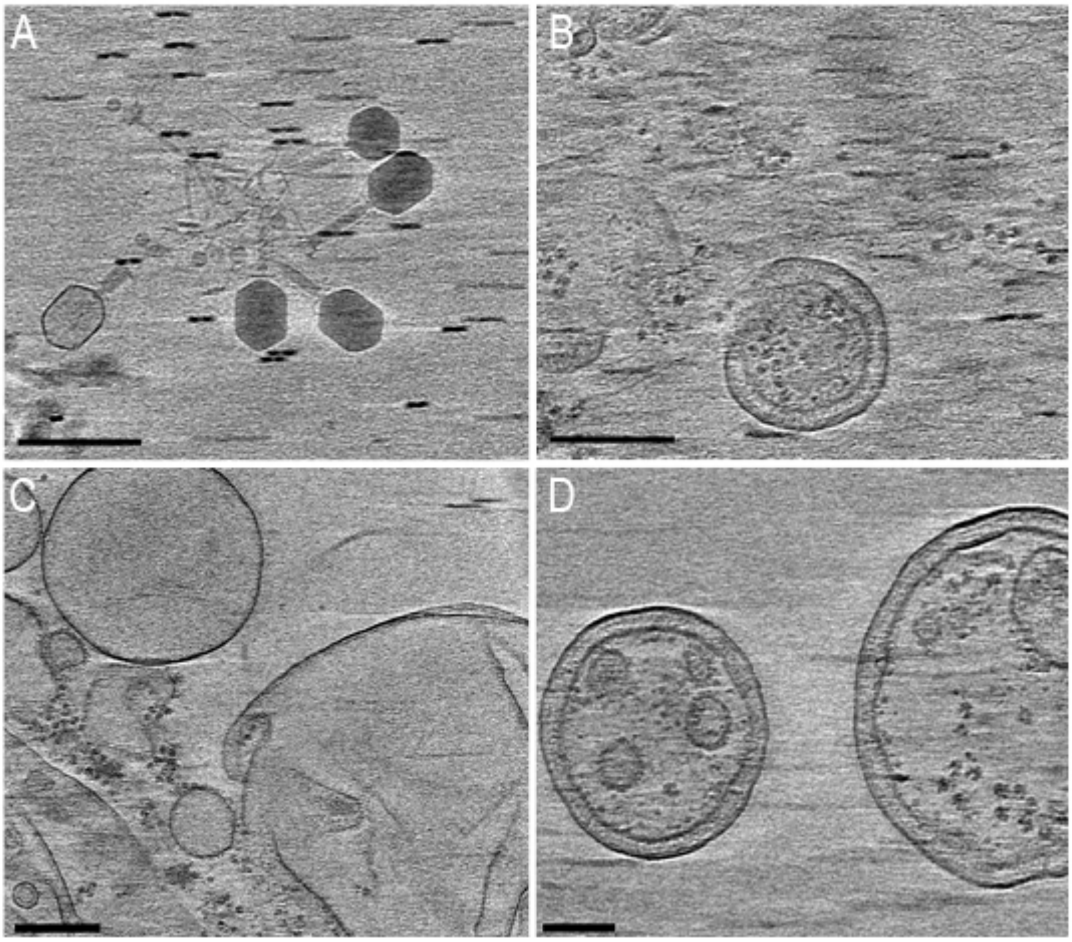


**Fig. S4. Heterogeneous Cryo-ET sample with T4 phages and *E. coli* spheroplasts remnants.** (**A**) Four unattached phages with a full capsid head and one with an empty capsid head. (**B**) Image of spheroplast fragments taken from the same tomogram as panel A. Phages were in close proximity to these fragments but not attached. (**C**) Typical view of the tomography data showing empty vesicles and outer membrane fragments. No attached phages were observed in any of the datasets. (**D**) Remnants of spheroplasts with both an inner and outer membrane, without phages in proximity. Scale bars represent 200 nm.

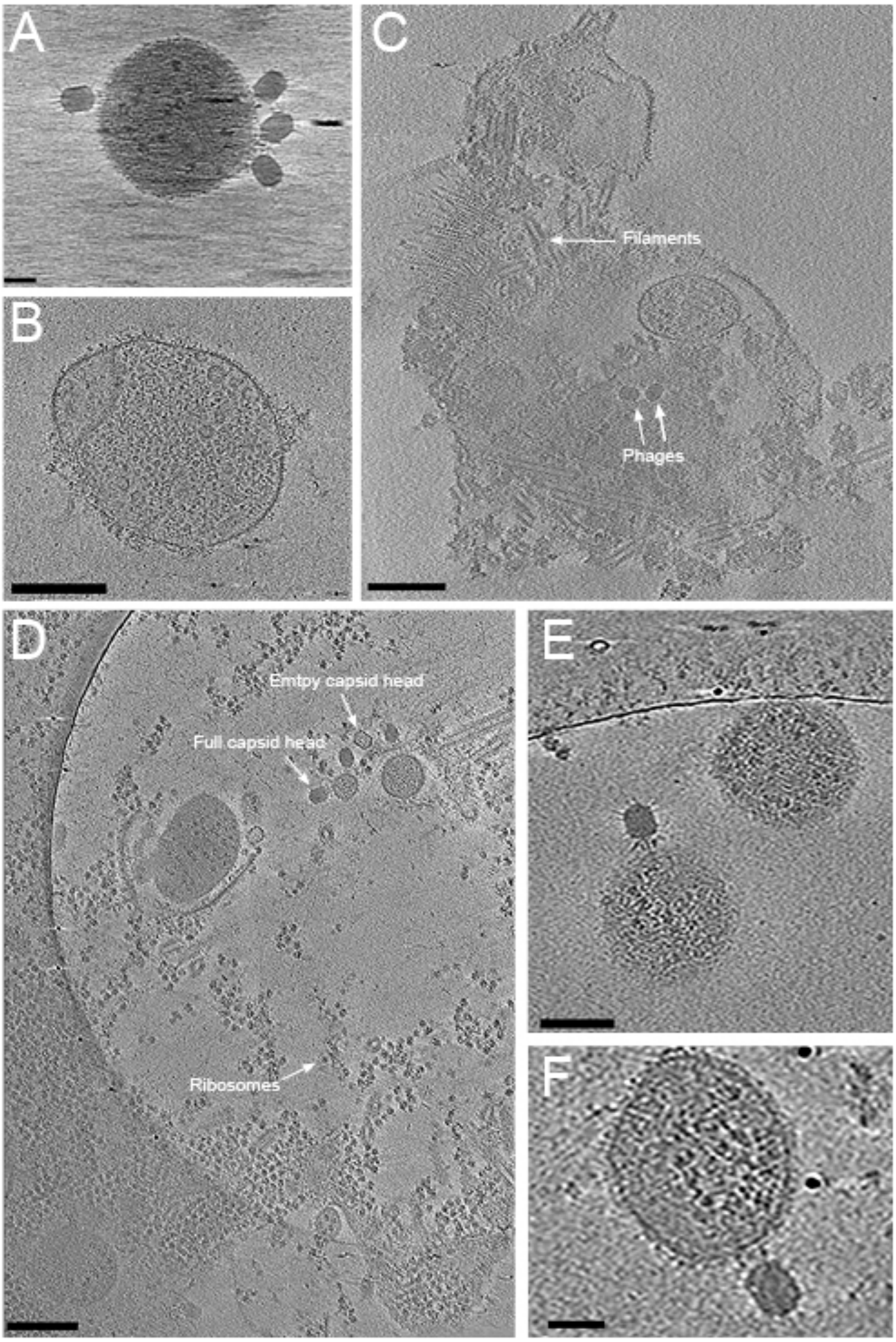


**Fig. S5. Φ29 Can attach to *B. subtilis* protoplasts and produce progeny.** (**A**) Four φ29 phages with a full capsid head attaching to a *B. subtilis* protoplasts. (**B**) A protoplast with ribosomes and filaments inside. (**C**) An exploding protoplast extruding phages, filaments and cell debris in the environment. (**D**) A protoplast exploding at the bottom and phages with full and empty capsid heads attaching to CWD cells at the top. (**E**) Phage φ29 with a retracted sheet in the process of ejecting DNA inside a protoplast. (**F**) Phage φ29 attaching to a protoplast. Scale bars represent 30 nm (panel A), 50 nm (panel E and F) and 200 nm (panel B, C and D).

**Table S1. Strains used in this study**

| Strains | Genotype | Reference |
|---|---|---|
| **Bacterial strains** | | |
| *E. coli* RP437 | *thr-1 araC14 leuB6(Am) fhuA31 lacY1 tsx-78 λ-eda-50 hisG4(Oc) rfbC1 rpsL136(strR) xylA5 mtl1 metF159(Am) thiE1*, with pBTOK-sfGFP | Lab collection |
| *E. coli* ET8 | Wildtype | Lab collection (*7*) |
| *Bacillus subtilis* | Type strain 110NA | Franklin Norbrega, DSMZ |
| *Streptomyces coelicolor* (A3)2 M145 | Wildtype | Lab collection |
| *Streptomyces albus* G. ATCC 25426 | Wildtype | Lab collection |
| *Streptomyces lividans* 1326 | Wildtype | Lab collection |
| *Streptomyces sp.* MBT13 | Wildtype | Lab collection (*7*) |
| *Streptomyces sp.* MBT61 | Wildtype | Lab collection (*7*) |
| *Streptomyces sp.* MBT66 | Wildtype | Lab collection (*8*) |
| *Streptomyces sp.* MBT86 | Wildtype | Lab collection (*7*) |
| **Phages** | | |
| T4 | Wildtype | Lab collection |
| Φ29 | Wildtype | Franklin Norbrega |
| Phage CE2 | Wildtype | This work |
| Phage CE3 | Wildtype | This work |
| Phage CE10 | Wildtype | This work |
| Phage GA3 | Wildtype | This work |
| Phage LA7 | Wildtype | This work |
| Phage LD10 | Wildtype | This work |

**Supplementary Movie Legends**

**Movie S1.** Time laps of MBT86 mycelium embedded in 0.5% DNB soft agar followed for 48 hours after LA7 infection. The mycelium is losing density over time and starts to produce CWD cells after approximately 12 hours. Images were acquired automatically every 30 minutes for at least 48 hours.

**Movie S2.** Time laps of *B. subtilis* embedded in 0.5% LB soft agar followed for 48 hours after φ29 infection. After 12 hours, all rod-shaped cells have switched to a spherical morphology. Images were acquired automatically every 30 minutes for at least 48 hours.

**Movie S3**. Time laps of green fluorescent *E. coli* embedded in 0.5% LB soft agar induced with 20ng/ml anhydrotetracycline followed for 48 hours after T4 infection. Most bacteria lyse within 6 hours, but some persistent CWD cells survive. Images were acquired automatically every 20 minutes for at least 48 hours.

**Movie S4**. Tomogram of *E. coli* spheroplasts with five phages that did not attach to the cells.

**Movie S5.** Tomogram of *B. subtilis* protoplasts filled with filaments associated with phage DNA assembly and φ29 attaching to the membrane with full capsid heads and on the bottom a phage with an empty capsid head.
